# Changes in dynamic transitions between integrated and segregated states underlie visual hallucinations in Parkinson’s disease

**DOI:** 10.1101/2021.06.21.449237

**Authors:** Angeliki Zarkali, Andrea I. Luppi, Emmanuel A. Stamatakis, Suzanne Reeves, Peter McColgan, Louise-Ann Leyland, Andrew J. Lees, Rimona S. Weil

## Abstract

**Background:** Visual hallucinations in Parkinsons disease (PD) are transient, suggesting a change in dynamic brain states. However, the causes underlying these dynamic brain changes are not known.

**Methods:** Focusing on fundamental network properties of integration and segregation, we used rsfMRI to examine alterations in temporal dynamics in PD patients with hallucinations (n=16) compared to those without hallucinations (n=75) and a group of normal controls (n=32). We used network control theory to examine how structural connectivity guides transitions between functional states. We then studied the brain regions most involved in these state transitions, and examined corresponding neurotransmitter density profiles and receptor gene expression in health.

**Results:** There were significantly altered temporal dynamics in PD with hallucinations, with an increased proportion of time spent in the Segregated state compared to non-hallucinators and controls; less between-state transitions; and increased dwell time in the Segregated state. The energy cost needed to transition from integrated-to-segregated state was lower in PD-hallucinators compared to non-hallucinators. This was primarily driven by subcortical and transmodal cortical brain regions, including the thalamus and default mode network regions. The regional energy needed to transition from integrated-to-segregated state was significantly correlated with regional neurotransmitter density and gene expression profiles for serotoninergic (including 5HT2A), GABAergic, noradrenergic and cholinergic but not dopaminergic density profiles.

**Conclusions:** We describe the patterns of temporal functional dynamics in PD-hallucinations, and link these with neurotransmitter systems involved in early sensory and complex visual processing. Our findings provide mechanistic insights into visual hallucinations in PD and highlighting potential therapeutic targets.

## Introduction

Visual hallucinations are a common symptom of Parkinson’s disease (PD) and are associated with cognitive decline(1), poorer quality of life(2) and increased mortality(3). The brain changes that give rise to hallucinations are not fully understood. However, the transient nature of hallucinations, even in patients who regularly experience them, suggests that the underlying process is a dynamic one and may be best examined using imaging techniques sensitive to dynamic changes in brain states. One potential approach is resting state functional MRI (rsfMRI), which measures spontaneous fluctuations in brain activity based on correlated fluctuations in blood oxygenation(4). This technique has shown changes in relative activity of specific functional brain networks in patients with PD-hallucinations(5), characterised by increased activation of the default mode network (DMN) and impaired recruitment of the dorsal attention network(6–9). However, these studies only provide a static image of functional connectivity, calculated over an entire scanning period, rather than examining dynamic changes in brain states.

An extension of this approach is dynamic functional connectivity analysis, which measures spontaneous fluctuations in connectivity over time(10–12) and may be a more accurate representation of fluctuating cognitive states than previous static approaches(13). Changes in temporal dynamics are seen in schizophrenia and other psychiatric conditions(14–17), and recent work showed imbalance of temporal dynamics of *integrated* and *Segregated* states in anaesthesia and disorders of consciousness(18,19) and after administration of the psychedelic LSD-25, known for its hallucinogenic properties(20). Changes in dynamic functional connectivity have been described in PD(21) and are associated with the severity of both motor and cognitive symptoms(22–24) but are as yet unexplored in relation to neuropsychiatric symptoms.

PD patients with hallucinations show widespread disruption in structural connections between brain regions, measured using diffusion MRI(25,26). These changes particularly affect highly connected brain regions or “hubs” important for switching the brain between different states(27,28). This is important because structural connectivity constrains the temporal alteration between different brain states(29,30) and the transitions between states can be modelled using network control theory(31). Specifically, the optimal energy cost needed to move the brain from one state to another can be calculated based on its structural network(31–33). A state that is less energy-demanding to maintain, or requires lower energy for transition, will be preferred. This framework can explain why a particular state is predominantly seen in health and how the balance between states may change in the presence of disease.

In this study we aimed to investigate the pattern of temporal dynamics in PD-associated visual hallucinations using rsfMRI; and determine whether the balance between integrated and Segregated states is preserved in patients with PD and visual hallucinations compared to those without hallucinations and controls. We found that patients with Parkinson’s hallucinations show impaired temporal dynamics, with a predisposition towards a Segregated state. We then used network control theory to calculate each individual’s required energy cost to transition from the integrated-to-the-Segregated state and vice versa, and the cost to maintain each state. We found that patients with Parkinson’s hallucinations required less energy to transition from the integrated-to-Segregated state than those without hallucinations and controls. Finally, we identified the brain regions that contribute most to transitions from integrated-to-Segregated state. As dynamic neural systems are, at least partly, modulated by neurotransmitter systems(34) we related the regional pattern of this transition to neurotransmitter systems using PET-derived regional neurotransmitter density profiles and regional gene expression for neurotransmitter receptors.

## Methods and Materials

### Participants

123 participants were included: 91 PD patients and 32 unaffected controls. All PD patients fulfilled Queen Square Brain Bank Criteria(35) and were recruited from clinics in the National Hospital for Neurology and Neurosurgery and affiliated hospitals. Controls were recruited from spouses and volunteer databases.

Patients with PD were classified as PD with visual hallucinations (PD-VH, n=16) if they scored ≥ 1 in Question 2.1 of the Unified Parkinson’s Disease Rating Scale (UPDRS). All other patients were classified as PD-non-VH (n=75). We collected additional information on severity, frequency and the phenomenology of experienced hallucinations with the University of Miami Parkinson’s Disease Hallucinations Questionnaire (UM-PDHQ)(36).

Study participants underwent clinical assessments of general and specific cognition across 5 domains as well as PD-specific measures (*Supplementary Methods 1*).

## Results

Study participants included 16 PD patients with habitual visual hallucinations (PD-VH), 75 PD patients without hallucinations (PD-non-VH) and 32 controls. PD-VH and PD-non-VH were well matched in demographics, cognitive and motor performance, levodopa equivalent dose, and image quality (Supplementary Table 1).

### Preserved topology of dynamic functional connectivity states

To examine the dynamic changes in functional connectivity underlying PD-hallucinations, we employed an a-priori clustering of dynamic functional connectivity into two states, an *Integrated* and a *Segregated* state. We found no significant differences within each state between PD versus controls or PD-VH versus PD-non-VH when comparing connectivity strength in each state using network-based statistics. We also found no significant between-group differences in the *Integrated* and *Segregated* states in terms of connectivity density (*Integrated*: Kruskal Wallis H=2.473, p=0.290, *Segregated*: 0.175, p=0.529), entropy of connectivity values (*Integrated*: H=0.723, p=0.696, *Segregated*: H=0.905, p=0.636), structural-functional coupling (*Integrated* F(111,2)=1.093, p=0.339, *Segregated*: F(111,2)=1.401, p=0.251) or small world propensity (*Integrated*: H=1.065, p=0.587, *Segregated*: H=4.400, p=0.111).

### Impaired temporal properties of dynamic functional connectivity in Parkinson’s hallucinations

Although the states themselves were preserved between groups in terms of network properties, we found significant changes in their temporal properties. PD-VH spent a significantly smaller proportion of time in the *Integrated* state (therefore higher proportion of time in the *Segregated* state) than PD-non-VH (β=-0.113, p=0.032) and than controls (β=-0.128, p=0.026) (Figure 2A). Within PD patients, the proportion of time spent in the *Integrated* state was inversely correlated with hallucination severity, measured by the UM-PDHQ (ρ=-0.259, p=0.013) (Figure 2B). Mean dwell time in the *Segregated* state was higher in PD-VH than PD-non-VH (19.1±16.9 in PD-VH vs 9.5 ± 9.1 in PD-non-VH H=4.058, p=0.044), with no difference between the two groups in mean dwell time of the *Integrated* state (H=2.166, p=0.141). No differences were seen in dwell time of either state between PD and controls. Finally, the total number of transitions was lower in PD-VH than PD-non-VH (5.7±5.3 in PD-VH vs 8.5±6.2 in PD-non-VH, H=3.87, p=0.049) but there was no difference between the two groups when the number of transitions from *integrated*-to-*Segregated* state and *Segregated*-to-*integrated* state were examined separately.

**Figure 1.**
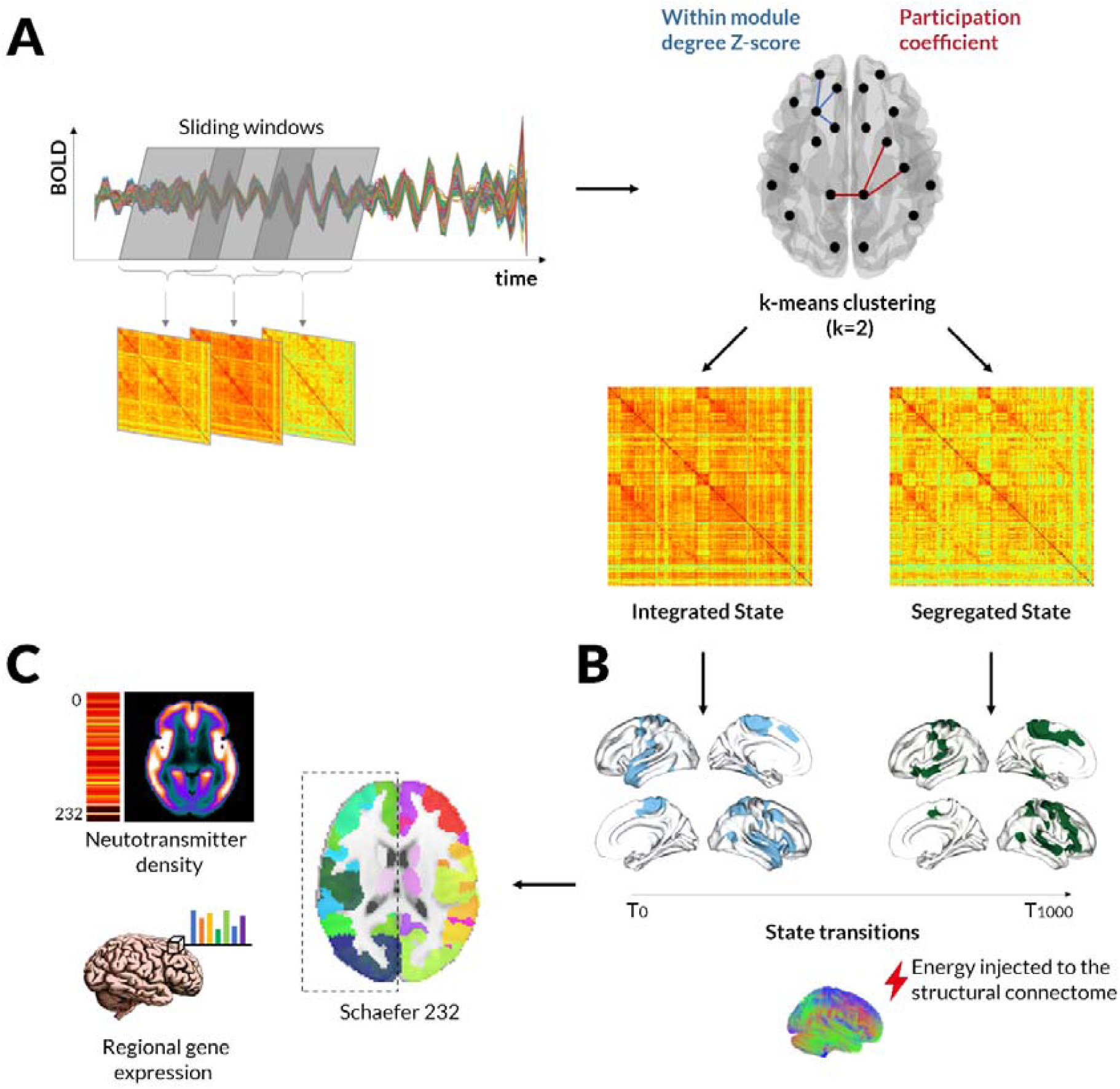
Overview of the study methodology. **A. Deriving integrated and segregated states of dynamic functional connectivity**. After obtaining sliding-windows (each 44s duration) of dynamic functional connectivity for each participant, the joint histogram of participation coefficient and within-module degree Z-score was used for k-means clustering (k = 2). (BOLD, blood oxygen level dependent activity). The cluster with highest average participation coefficient is then identified as the predominantly *Integrated* dynamic state and the cluster with the lowest participation coefficient as the predominantly *Segregated* state. Note that this is done for each participant separately leading to individually-defined *Integrated* and *Segregated* states. **B. Modelling state transitions**. After deriving each individual’s *Integrated* and *Segregated* states we used an optical control framework to calculate the minimal control energy that needs to be applied to each node of the structural network to transition from a baseline state at time T0 to a target state at time T1000. Here, as an example, we illustrate the transition from the *Integrated* state (top 20% of nodes in blue) to the *Segregated* state (top 20% of nodes in green) but minimal energies were also calculated for *Segregated*-to-*Integrated* transition as well as minimal energies to maintain the *integrated* state (*Integrated*-to-*integrated*) and *Segregated* state (*Segregated*-to-*Segregated*) using the same model. Minimal control energies were calculated for each subject based on their structural brain network, which was estimated using diffusion imaging and probabilistic tractography. Both states were represented in the model as a vector of the sum connectivity strength for each node (1*232). **C. Linking with neurotransmitter systems**. Minimal control energies to transition between and maintain functional states were compared between patients with PD with (PD-VH) and without hallucinations (PD-non-VH). Transitions that differed between groups were then further explored to examine whether contributing nodes (requiring mode control energy) were associated with specific neurotransmitter systems. To do this, we calculated for each of the 232 regions of interest of our parcellation (Schaeffer 232: 200 cortical and 32 subcortical regions) 1) mean neurotransmitter density profiles derived from PET data (serotonin (5HT1a, 5HT2a and 5HT1b), dopamine (D1 and D2) and GABAA receptors) and 2) gene expression profiles for each of 31 pre-selected genes encoding receptors for norepinephrine, acetylcholine, dopamine and serotonin.

**Figure 2.**
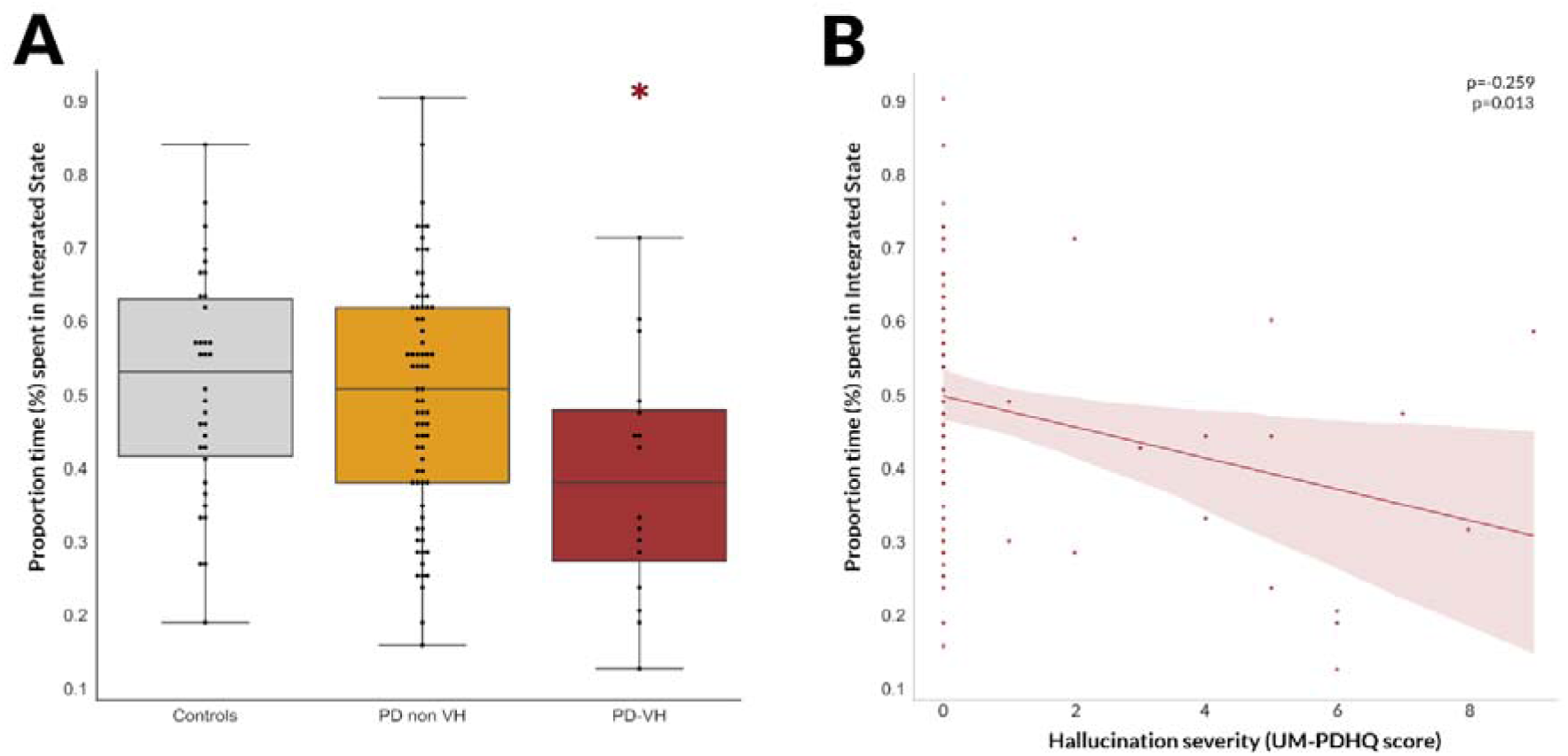
Altered temporal properties of dynamic functional connectivity in patients with Parkinson’s and visual hallucinations. **A. Percentage of total time spent in the Integrated state**. Patients with Parkinson’s with visual hallucinations spent significantly less time in the *Integrated* state of dynamic functional connectivity than patients without hallucinations (p=0.032) and controls (p=0.0262). (Error bars are 95% confidence intervals) **B. Correlation between proportion of time spent in the Integrated state and hallucination severity** The proportion of time spent in the *integrated* State was significantly correlated with hallucination severity in participants with Parkinson’s disease (Spearman’s correlation coefficient ρ=-0.259, p=0.013): participants with more severe hallucinations spent less time in the *integrated* state. PD-VH: Parkinson’s disease with visual hallucinations, PD-non-VH: Parkinson’s disease without hallucinations. UM-PDHQ: University of Miami Parkinson’s disease Hallucinations Questionnaire, higher scores indicate more severe and frequent hallucinations.

Overall, this suggests that PD-VH spend more time in the *Segregated* state than PD-non-VH, with fewer total transitions and longer dwelling time within the *Segregated* state (Figure 2).

### Reduced energy costs to transition from the integrated to segregated state in Parkinson’s patients with visual hallucinations

Having identified significant differences in terms of brain dynamics between PD-VH and PD-non-VH, which are specifically related to the severity of visual hallucinations (the focus of our present investigation), we sought to interrogate further this difference between PD patients. Specifically, we aimed to investigate whether the *Segregated* state predominance observed in PD-VH participants could be explained by differences in ease of transition from the *integrated*-to-*Segregated* state or vice versa or a difference in ease of maintaining the *Segregated* state in PD-VH compared to PD-non-VH participants. To do this, we calculated the minimal control energy that needs to be applied to the structural network of each participant to 1) transition from *integrated*-to-*Segregated* state 2) transition from *Segregated*-to-*integrated* state 3) maintain the *integrated* state and 4) maintain the *Segregated* state. We then examined whether transition and persistence energies in each state differed between PD-VH and PD-non-VH.

Similarly to previous work(64), persistence energy for the computationally more demanding *Integrated* state was higher than the *Segregated* state for all participants (log(Persistence Energy) *Integrated*: 13.8±0.9 vs 13.7±1.1 *Segregated*, repeated measures ANOVA main effect of *Integrated* to *Segregated* state persistence energy F(1,113)=12.432, p<0.001). Similarly the minimal energy needed to transition from the less connected *Segregated* to the more interconnected *Integrated* state was higher (*Integrated*-to-*Segregated* 13.9±0.7 vs *Segregated*-to-*Integrated* 14.1±0.8, F(1,113)=6.722, p=0.011) (Supplementary Figure 2).

PD-VH needed significantly lower control energy to transition from the *integrated*-to-*Segregated* state than PD-non-VH (effect size Hedge’s g=0.922, t=2.376, p=0.029) (Figure 3A). The minimal control energy to transition from an *Integrated*-to-*Segregated* state was significantly correlated with hallucinations severity: lower energy associated with more severe hallucinations (ρ=-0.283, p=0.008) (Figure 3B). There were no statistically significant differences between PD-VH and PD-non-VH in the minimal control energy needed to transition from *Segregated*-to-*integrated* state (t=1.346, p=0.195), or to persist within the *Integrated* (t=1.041, p=0.312) or *Segregated* state (t=1.079, p=0.295). Therefore, network control theory reveals that the higher proportion of time that PD-VH patients spend in the *Segregated* state may be accounted for in terms of this state being easier to transition to from the *Integrated* state (as opposed to being easier to persist in).

**Figure 3.**
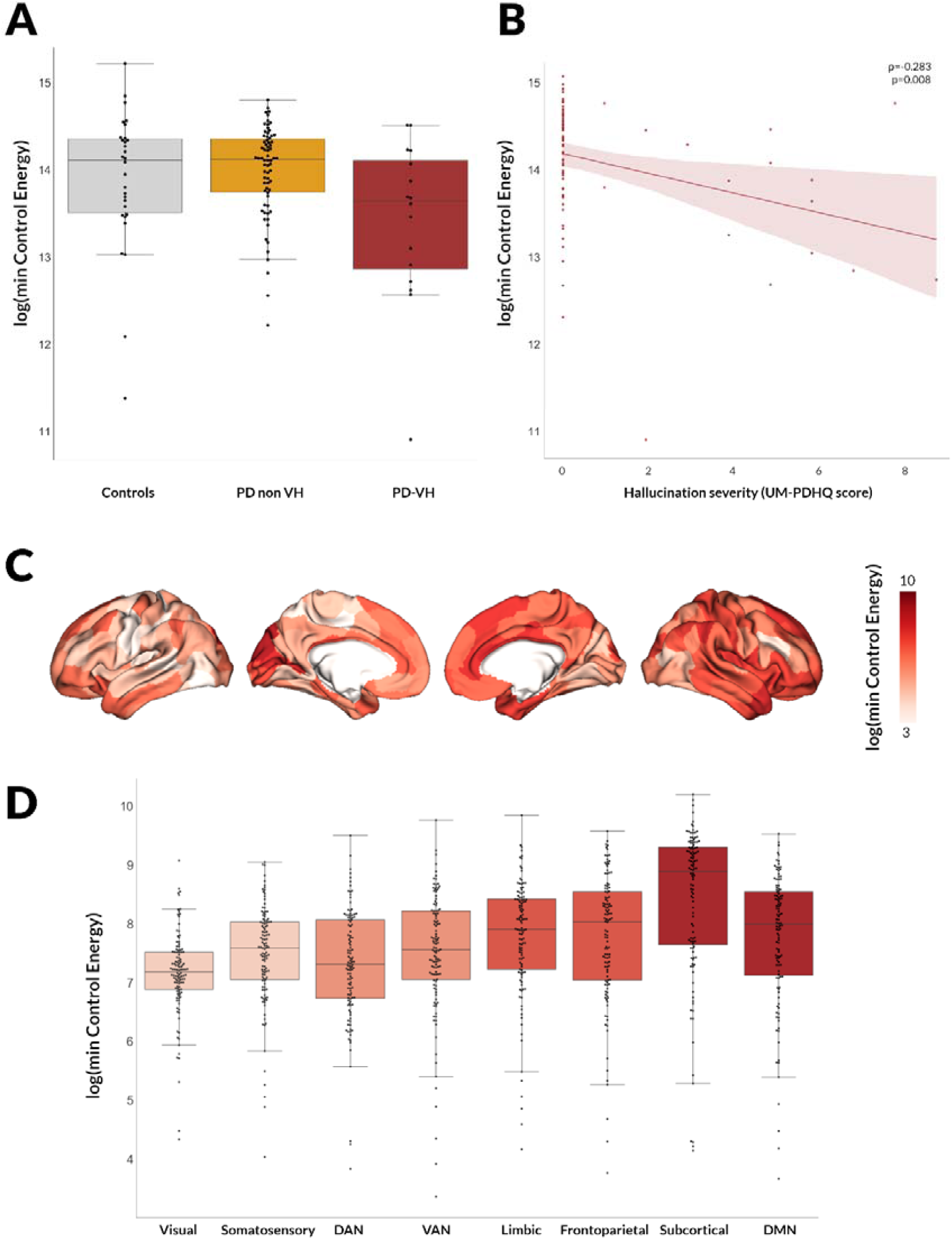
Changes in control energy to transition from the Integrated to the Segregated state in patients with Parkinson’s and visual hallucinations. **A. Minimal control energy to transition from the Integrated to the Segregated state** Less energy is needed to transition for patients with Parkinson’s and visual hallucinations (PD-VH) than those without hallucinations (PD-non-VH). Log-transformed minimal control energy is presented. Error bars are 95% confidence intervals **B. Correlation between minimal control energy and hallucination severity** The log-transformed minimal control energy required across the whole of the network to transition from the *Integrated* to the *Segregated* state was significantly correlated with severity in participants with Parkinson’s disease (Spearman’s correlation coefficient ρ=-0.283, p=0.008): participants with more severe hallucinations needed less energy to transition, suggesting that the transition to the *Segregated* state may be easier to achieve in PD-VH therefore prefered. **C. Regional variation in minimal control energy to transition from the Integrated to the *Segregated* state** The log-transformed minimal control energy that needs to be applied to each node is presented; darker colours denote higher amounts of energy required. Note that only cortical regions are plotted. **D. Minimal control energy per functional subnetwork**. The mean minimal control energy to transition from the Integrated to the Segregated state across all nodes of the seven cortical and one subcortical resting state networks is plotted. Darker colours denote higher levels of the cortical hierarchy; also left to right: unimodal to transmodal regions. There was a significant correlation between the minimal transition energy from integrated-to-Segregated state that was needed to be applied to each node and the nodes position in the cortical hierarchy, with higher amount of energy needed for more transmodal regions (ρ= 0.526, p<0.001).

**Figure 4.**
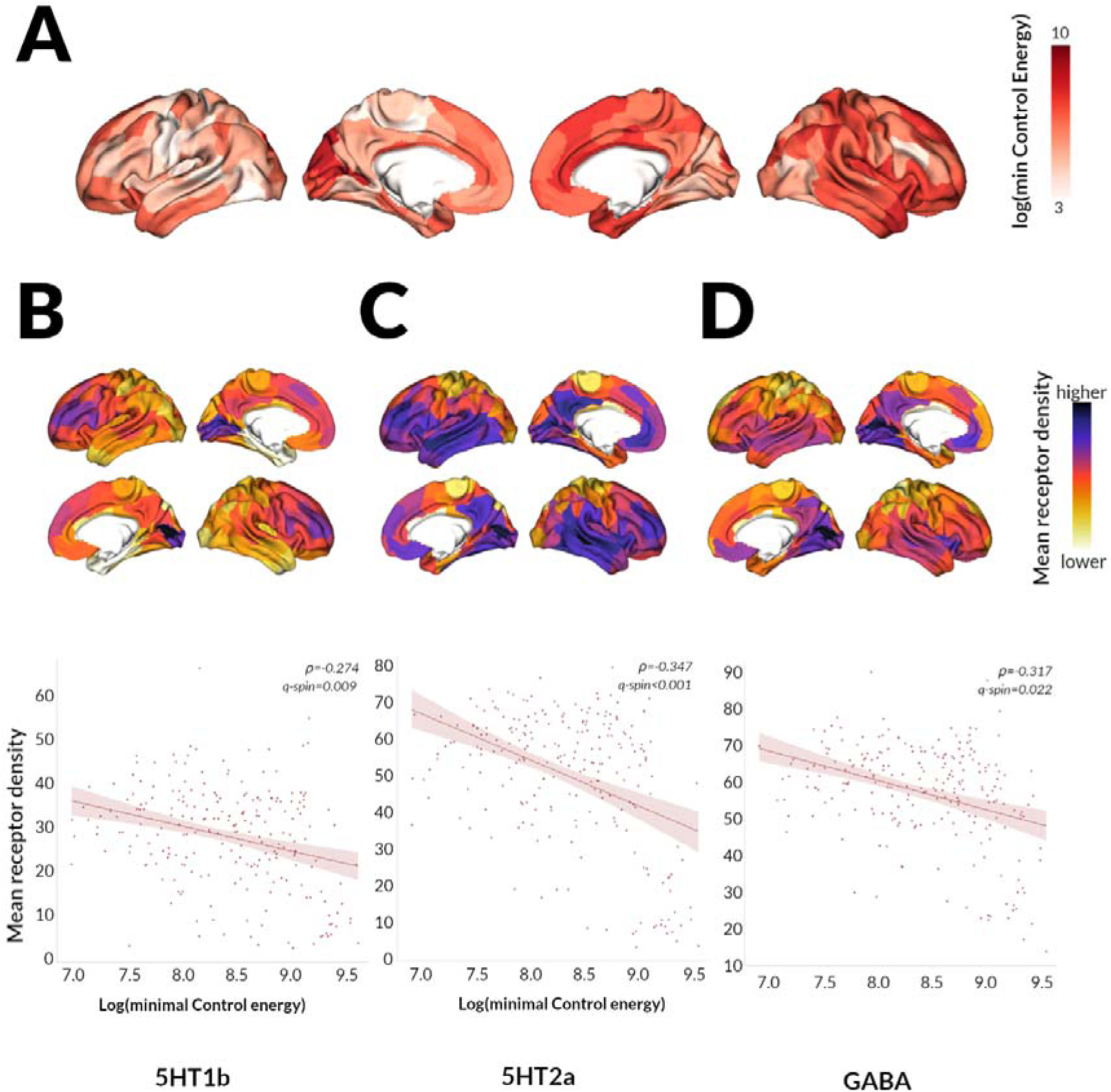
Neurotransmitter correlates of Integrated-to-Segregated state transition. The log-transformed minimal control energy that needs to be applied to each node (A) was correlated with the mean regional receptor density of 5HT1b receptors (B), 5HT2a receptors (C) and GABA receptors (D), from open access atlases of PET data in unaffected individuals. In all cases ρ is the Spearman correlation coefficient and q-spin is the FDR corrected p-value derived following spatial permutations (p-spin, 1000 permutations).

### Transition from integrated to the segregated state is driven by subcortical and more multimodal brain regions

Next, we aimed to identify which brain regions contribute more to the transition from the *Integrated*-to-*Segregated* state (which nodes need more energy to be applied to transition). As expected(65), subcortical regions heavily contributed, with 25 subcortical nodes amongst the top 20% of contributors (25/47 or 53.2%) with thalamic regions amongst the highest contributors. Amongst cortical nodes, top contributors included primarily right hemispheric regions (20/22 cortical nodes) including regions of the Default mode network: cingulum, precuneus, inferior and superior temporal regions and medial frontal regions (Figure 3C). There was a significant correlation between the transition energy from *Integrated*-to-*Segregated* state that needed to be applied to each node and the node’s position in the cortical hierarchy, with higher energy needed for more transmodal regions (ρ= 0.526, p<0.001) (Figure 3D for regional energy per functional network).

### Correlation with neurotransmitter systems

Finally, we examined whether the *Integrated*-to-*Segregated* state transition, which was less costly for PD-VH patients is associated with specific neurotransmitter systems in the healthy brain. To do this, we correlated the mean control per node to transition from the *Integrated*-to-*Segregated* state with mean regional neurotransmitter density (derived from open-access PET data) and neurotransmitter receptor gene expression levels (derived from the Allen Brain atlas(61)) in health; we tested this against spatially-correlated null models through sphere permutations, FDR-corrected for multiple comparisons.

We found a significant correlation between regional log(Energy) and density of 5-HT1b (ρ=-0.274, q_spin_=0.009), 5-HT2a (ρ=-0.347, q_spin_<0.001) and GABAA receptors (ρ=-0.317, q_spin_=0.022), from open-access atlases of PET data. Regional energy and regional expression levels of genes relating to 5-HT2a receptors were also significantly correlated (ρ=-0.1438, q_spin_=0.044) as well as two GABA_A_ receptors [GABRA1 (ρ=-0.2437, q_spin_=0.020) and GABRA2 (ρ=0.128, q_spin_=0.023)]; gene expression data for 5-HT1b receptors were not available. Although noradrenergic and acetylcholinergic PET data are not publically available, genetic expression of noradrenergic (ADRA1B and ADRA2A), muscarinic (CHRM1, CHRM2, CHRM3, CHRM4) and nicotinic receptors (CHRNA3, CHRNA4, CHRNA7, CHRNB2) was correlated with regional transition energy. Gene expression of DRD2 was also correlated with regional control energy for the *Integrated*-to-*Segregated* state transition (ρ=0.318, q_spin_=0.013) but this was not replicated using density PET-derived data (ρ=0.056, q_spin_=0.800).

## Discussion

We have used dynamic functional connectivity and network control theory to explore the temporal dynamics underlying visual hallucinations in Parkinson’s, and examine how these specific patterns of temporal dynamics can be explained through brain structure. We found that PD-hallucinators spent more time in a *Segregated* state of functional connectivity than those without hallucinations, with fewer total transitions and longer dwelling time within the *Segregated* state. The transition from the *Integrated*-to-*Segregated* state was less energy demanding in PD-hallucinators than non-hallucinators. This transition is mediated by trans-modal brain regions that are associated with specific neurotransmitter systems, as confirmed through combined in-vivo PET mapping and post-mortem gene expression microarray data.

Previous studies have shown that PD patients with cognitive impairment similarly spend more time in a Segregated state and show fewer transitions between states than PD with intact cognition and controls(23,24). There were no differences in cognitive performance between PD patients with and without hallucinations in our cohort, but visual hallucinations are known to be associated with incipient dementia in PD(66).

In schizophrenia, where auditory hallucinations are a core feature, similar findings of altered dwell time are seen(14,67), correlated with severity of hallucinations(68). We similarly saw a correlation with hallucination severity with patients with more severe visual hallucinations spending less time in the *Integrated* (and more time in the *Segregated*) state suggesting this finding is specific to hallucinations as a trait.

Only the temporal dynamics of functional connectivity were altered in PD with hallucinations. This indicates that a change in the temporal balance between normal/preserved states rather than a change in the states themselves underlie PD-hallucinations. Using similar methodologies, studies in loss of consciousness and after LSD-25 ingestion in healthy volunteers show within-state changes particularly within the *Integrated* state(18,20), which is more linked to cognitive performance and alertness(12). As we examined the propensity to hallucinate rather than the hallucinatory state (participants were not actively experiencing hallucinations during scanning) it is possible that additional within-state changes could underlie visual hallucinations in PD, in the moment when they actually occur, an avenue for potential future investigations. In addition, although hallucinations in our participants were frequent (at least weekly) they were not universally complex and severe. In contrast, the acute LSD-induced visual hallucinations are believed to be due to serotoninergic system activation alone (specifically the 2A receptor(69)), and are associated with changes in other sensory modalities including time/space dysperceptions and ego dissolution(70), which are not seen with PD-associated hallucinations; thus it is not unexpected that the underlying changes in temporal dynamics are different.

As temporal transition between functional states is constrained by structural connectivity(29,30,32), we examined the energy cost of transitioning between and maintaining the *Integrated* and *Segregated* states. There was a significantly lower energy cost to transition from the *Integrated*-to-*Segregated* state for PD-hallucinators than non-hallucinators, suggesting that this transition is easier to achieve in hallucinators. This transition is mediated primarily by subcortical and multimodal brain regions of DMN, further highlighting its involvement in PD hallucinations(6). Thalamic regions were amongst the highest contributors to this transition. Thalamic involvement has been previously described in visual hallucinations(56,71) and we recently showed longitudinal changes in grey and white matter within the medial mediodorsal thalamus(72). This provides further evidence of the thalamus as a key driver of network imbalance in PD-hallucinations(65,73).

Interestingly, the brain regions contributing most to this transition from *Integrated*-to-*Segregated* state showed a correlation with specific neurotransmitter systems in health. Although the directionality of the relationship is difficult to interpret as data on regional neurotransmitter density and gene expression were derived from healthy individuals, regional density of 5HT2A receptors was significantly correlated with the regional control energy needed for *Integrated*-to-*Segregated* state transition; this was replicated using regional expression data for the 5HT2A receptor gene.

Activation of 5HT2A receptors is a key mechanism for drug-induced hallucinations occurring with the psychedelic drugs, LSD-25, psilocybin and ayahuasca(74) and modelling studies have shown that this receptor plays a key role in engendering the characteristic brain dynamics of LSD(69). 5HT2A has also been implicated in PD-hallucinations; evidenced by the higher density of 5HT2A receptors within frontal, temporal and occipital regions in patients with PD hallucinations in post mortem and in vivo studies(75,76) and the efficacy of the novel 5HT2A inverse agonist Pimavanserin in the treatment of PD-hallucinations(77).

Other serotonergic receptors were also important for the *Integrated*-to-*Segregated* state transition including: 5HT1B (receptor density, no genetic expression data), 5HT1E, 5HT1F and 5HT5A (gene expression data only). The correlation with multiple serotonin receptors, indicates that serotonergic modulators targeting multiple receptors could be potential therapeutic targets for PD-hallucinations. Of note, no receptor density or gene expression data were available for 5HT3 receptors, a target of interest for Ondansetron, 5HT3-antagonist currently under evaluation as a treatment of hallucinations(78).

Regional receptor density and gene expression for GABAergic receptors were also correlated with regional transition energy in line with previous studies showing reduced GABA concentration in the visual cortex of PD-hallucinators(79,80). Visual processing involves a complex interplay between monoaminergic, cholinergic and GABA/glutamatergic neurotransmission(73). The observed correlation between the *Integrated*-to-*Segregated* state transition and regional gene expression of noradrenergic (ADRA1B, ADRA2A) and cholinergic (muscarinic and nicotinic) receptors is consistent with this, but there was no available PET derived density data to replicate this.

Convergent evidence has recently highlighted the importance of the noradrenergic system in some non-motor PD symptoms(81–83). Noradrenaline plays a key role in modulating selective attention(84) and with serotonin, modulates behavioural responses to incoming visual information(73). Changes within the noradrenergic system may be involved in altered state transitions in PD-hallucinations by modulating the activity of sensory cortices and thalamocortical neurocircuitry(85). In contrast there was no consistent correlation with dopaminergic receptors. These findings highlight the role of transmitters other than dopamine in the development of PD-hallucinations.

Several considerations need to be taken into account when interpreting our findings. Functional data are susceptible to motion artefact; we adopted strict exclusion criteria to mitigate for this(86). Global signal regression is a potential additional tool to counteract residual artifacts from head motion(86) however it may contain behaviourally-relevant information and affect group results(40,87), therefore we did not regress global signal (18,20). All participants were scanned while receiving their usual dopaminergic medications and at the same time of day and levodopa equivalent doses did not significantly differ between PD-VH and PD-non-VH(88). Further studies assessing PD patients ON and OFF levodopa might provide additional information. Although brain networks are non-linear, we used a linear optimal control model since this has been shown to provide important insights on non-linear dynamics(89) and linear-Gaussian models are often adequate descriptors of functional MRI timeseries, such that more complex, non-linear models often do not provide additional explanatory power(90,91). Nevertheless, future work may seek to leverage insights from non-linear models of brain dynamics, e.g. through neurobiologically detailed dynamic mean-field models that have already been successfully applied to the study of altered states of consciousness(69,92). Finally, data on neurotransmitter density and gene expression were not derived from our participants but from separate cohorts of healthy volunteers and post-mortem human brains respectively; therefore results relating to neurotransmitter receptors should be interpreted with caution. Future work may seek to replicate these results with each patient’s own unique neurotransmitter receptor signature, which may offer individualised insights and the opportunity to assess the directionality of this relationship, as well as potential targets for pharmacological intervention.

## Conclusions

Our findings describe, for the first time, that temporal functional dynamics are altered in PD-hallucinations, with a predisposition towards a Segregated state of functional connectivity. This Segregated state predominance can be explained by a reduced energy cost to transition from the integrated-to-Segregated state in PD patients with hallucinations compared to those without hallucinations. We have also clarified the neuromodulatory correlates of the integrated-to-Segregated state transition in the healthy brain. These results provide mechanistic insights into visual hallucinations in PD and possible therapeutic targets.

## Notes

### Competing Interest Statement

The authors have declared no competing interest.

